# The Serine Shunt enables formate conversion to formaldehyde *in vivo*

**DOI:** 10.1101/2024.07.31.605843

**Authors:** Karin Schann, Sebastian Wenk

## Abstract

Microbial valorization of CO_2_-derived substrates has emerged as a promising approach to address climate change and resource scarcity. Formate, which can be efficiently produced from CO_2_, shows great potential as a sustainable feedstock for biotechnological production. However, the scope of formate assimilation pathways is restricted by the limited number of natural formate-assimilating enzymes. To overcome this limitation, several new-to-nature routes for formate assimilation based on its reduction to formaldehyde have been proposed, but they suffer from low catalytic efficiencies and cannot yet support bacterial growth. Here, we propose the Serine Shunt as a novel formate reduction route and demonstrate its activity *in vivo*. In this pathway, formate is attached to glycine to form serine, which is subsequently cleaved into formaldehyde and glycine, thereby effectively converting formate to formaldehyde. Unlike other formate reduction routes, the Serine Shunt mainly utilizes natural reactions with favorable enzyme kinetics, while requiring the same amount of ATP and NAD(P)H as the most efficient new-to-nature route. We implemented the Serine Shunt in engineered *E. coli* strains using a step-wise approach by dividing the pathway into metabolic modules. After validating the individual module activities, we demonstrated the *in vivo* activity of the complete Serine Shunt by measuring intracellular formaldehyde production with a GFP sensor and coupling its activity to cell growth. Our results indicate that the Serine Shunt could be applied as a novel formate reduction route in methylotrophic hosts relevant for biotechnology.

## Introduction

The rapidly progressing climate change requires immediate actions to reduce the CO_2_ concentration in the atmosphere (Lee et al., 2022). This should be addressed by reducing CO_2_ emissions and by capturing CO_2_ from the atmosphere, followed by its storage or valorization. Biotechnological valorization of captured CO_2_ is a promising alternative to long-term storage as it recycles CO_2_ and minimizes the need for fossil or plant-derived products (Bachleitner et al., 2023). Here, reduced one-carbon (C1) molecules like formate or methanol could become important mediators as they can be electrochemically produced from CO_2_ and present favorable characteristics for microbial cultivation. In recent years, the field of C1 biotechnology has emerged as a subdiscipline which aims to combine chemical CO_2_ reduction with biotechnological conversion of C1 molecules to value-added products (Bachleitner et al., 2023; Orsi et al., 2023). The model of a formate-based bioeconomy proposes to explore formate as a favorable C1 substrate for the sustainable biotechnology (Yishai et al., 2016)

In nature, several bacteria grow aerobically on formate via the Serine Cycle (Anthony, 2011). Here, formate is assimilated into methylene-THF (CH_2_-THF) which is condensed with glycine to form serine. Thereafter, a chain of reactions leads to the recycling of glycine and the production of glyoxylate as a biomass precursor. Based on the formate assimilation module of the Serine Cycle, several new-to-nature metabolic pathways for formate assimilation in model microbes were suggested (Bar-Even et al., 2013). In recent years, two of these pathways, the Reductive Glycine Pathway and the Serine Threonine Cycle have been successfully engineered in model microbes (Bang et al., 2020; Claassens et al., 2020; Kim et al., 2020; Turlin et al., 2022; Wenk et al., 2022). However, the growth characteristics of the engineered microbes is still suboptimal for biotechnological applications. Thus, growth on formate via these pathways should be improved, or other formate assimilation routes should be explored.

To increase the number of formate assimilation routes, formate could be converted to formaldehyde which is a more reduced C1 mediator with higher energy density. Furthermore, it is more reactive than formate and can be easily assimilated into metabolism. It has the same reduction level as biomass and many formaldehyde assimilation reactions exist (both natural and synthetic) (Bar-Even, 2016). For example, many methylotrophic bacteria and yeasts, relevant for biotechnology, grow on methanol-derived formaldehyde via the Ribulose Monophosphate Cycle and the Xylulose Monophosphate Cycle (Chistoserdova et al., 2009; Pham et al., 2022; Wefelmeier et al., 2023; Wegat et al., 2022). Furthermore, several promising new-to-nature pathways for formaldehyde assimilation such as the Homoserine Cycle, the Formolase Pathway, the SACA Pathway and the Erythrulose Monophosphate Cycle have been proposed (He et al., 2020; Lu et al., 2019; Siegel et al., 2015; Wu et al., 2023).

As a metabolic pathway that converts formate to formaldehyde does not seem to exist in nature, two new-to-nature enzymatic cascades were previously proposed: (I) formate activation to formyl-CoA or (II) formate activation to formyl phosphate followed by reduction of the activated intermediate to formaldehyde (Bar-Even, 2016). Although enzyme activities of both cascades could be demonstrated, they were not sufficient to support cellular growth, suggesting that further enzyme optimization is required (Hu et al., 2022; Nattermann et al., 2023; Wang et al., 2021).

In this work, we propose the Serine Shunt as an alternative route for formate conversion to formaldehyde (**Figure 1**). By leveraging metabolic reactions from different C1 assimilation pathways, we designed a linear route that assimilates formate into serine and releases formaldehyde via serine cleavage. Like in the natural Serine Cycle, formate is assimilated into CH_2_-THF which is attached to glycine to form the stable intermediate serine. Thereafter, the serine aldolase from the synthetic Homoserine Cycle is used to cleave serine into formaldehyde and glycine.

**Figure 1:**
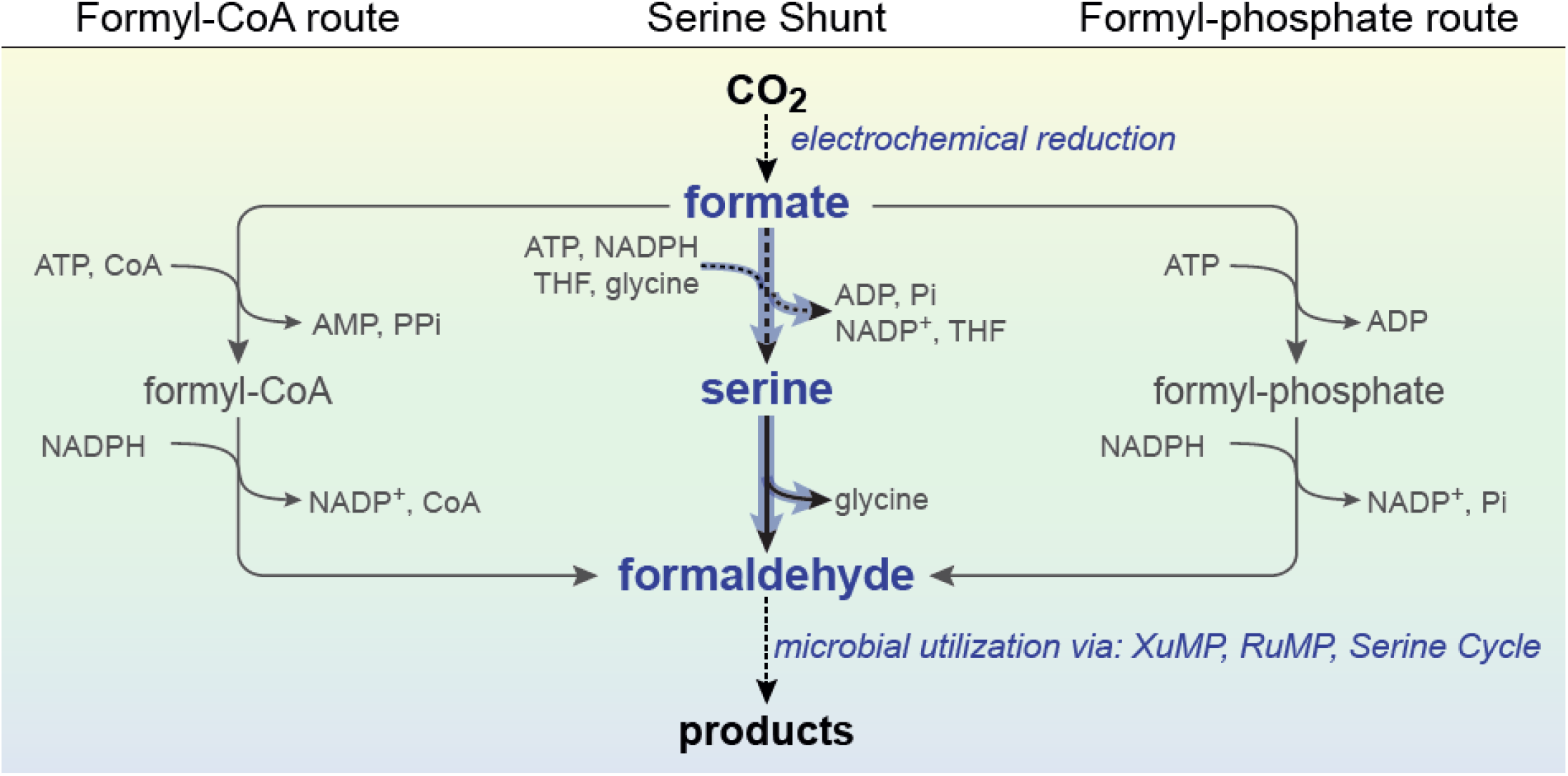
The Serine Shunt compared to other new-to-nature formate reduction routes. Reactions schemes for the different proposed formate reduction routes. The new-to-nature routes are shown in gray color. The Serine Shunt is shown in blue. Dotted arrows indicate multi-enzymatic reactions. Formate can be reduced into formaldehyde via two new-to-nature reaction routes, which use formyl-CoA of formyl-phosphate as intermediates. In the Serine Shunt formate is reduced to formaldehyde via its condensation with glycine to serine followed by serine cleavage to glycine and formaldehyde. The Serine shunt and the formyl phosphate route are the most energy efficient formate reduction routes as they consume one ATP and one NADPH to reduce one mole formate.

Using a step-wise engineering approach, we divided the Serine Shunt into two metabolic modules and tested the individual module activities in dedicated selection strains. Then, we combined both modules into a strain that depends on formaldehyde assimilation for cell growth and were able to show formate-dependent growth via the novel route. Using ^13^C-labelling we confirmed the conversion of formate to formaldehyde. The Serine Shunt presents a promising route to produce formaldehyde *in vivo* and could be directly applied in metabolic engineering studies aiming to establish novel formate assimilation pathways.

## Materials and methods

### Chemicals and reagents

Primers were synthesized at Integrated DNA Technologies (IDT). DNA fragments were amplified with the high-fidelity polymerase PrimeSTAR Max DNA Polymerase from TaKaRa and colony PCR reactions were performed using DreamTaq polymerase (Thermo Fisher Scientific). FastDigest restriction enzymes and T4 DNA ligase from Thermo Fisher Scientific were used for restriction-based cloning. The kits used for plasmid isolation and PCR cleanup were purchased from by Thermo Fischer Scientific. All chemicals used in the plate reader experiments were ordered from Sigma-Aldrich.

### Growth media

For cloning and strain engineering, cultures were grown in lysogeny broth (LB) medium (10 g/L NaCl, 10 g/L Tryptone, 5 g/L Yeast Extract) supplemented with the relevant antibiotic (50 μg/mL kanamycin or 100 μg/mL streptomycin). Diaminopimelate (DAP, 0.25 mM) needs to be supplemented to the growth media of the formaldehyde sensor due to the *asd* knockout. During growth experiments, cells were cultivated in M9 minimal medium (50 mM Na_2_HPO_4_, 20 mM KH_2_PO_4_, 1 mM NaCl, 20 mM NH_4_Cl, 2 mM MgSO_4_ and 100 μM CaCl_2_), supplemented with trace elements (134 μM EDTA, 31 μM FeCl_3_, 6.2 μM ZnCl_2_, 0.76 μM CuCl_2_, 0.42 μM CoCl_2_, 1.62 μM H_3_BO_3_, 0.081 μM MnCl_2_) and the relevant carbon sources (See paragraph **Growth experiments**).

### Strains

Plasmid construction was done in *E. coli* DH5α. The *E. coli* SIJ488 strain, which is derived from K-12 MG1655, was used as a base strain (Jensen et al., 2015). The KEIO collection was used as donor strains to prepare the P1 donor lysates for phage transduction (Baba et al., 2006).

### Strain engineering

The sensor strains were constructed by using recombineering techniques to modify the SIJ488 base strain. Deletions were made by P1 phage transduction as detailed in (Wenk et al., 2018). For P1 phage transduction, the KEIO collection strain with the desired knockout was grown in LB supplemented with 5 mM CaCl_2_ until an OD ∼0.3. Then the culture was infected with P1vir (Ikeda & Tomizawa, 1965) stock lysate for 2-4 hours. The cells were pelleted by centrifugation and the supernatant was filtered through a 0.22-μm filter to obtain the donor lysate that was then used for the transduction. The deletion in the relevant strain was made by resuspending the pellet of an overnight culture in P1 salts (LB supplemented with 10mM CaCl_2_ and 5mM MgSO_4_). P1 donor lysate was added to this cell suspension and incubated for 30 min at 30 °C. After a recovery for 1 hour in LB with ∼100 mM sodium citrate, the cells were plates on selective plates containing 5 mM sodium citrate and kanamycin. The antibiotics cassette was later removed by inducing the flippase. Knockouts and removal of the antibiotic cassette were confirmed by a colony PCR.

Genomic integrations were made using the recombineering machinery of the SIJ488 strain (Jensen et al., 2015). Therefore, the gene of interest was cloned together with a kanamycin antibiotic cassette, flanked by FRT sites, and amplified with primers containing 50 bp homology arms that are flanking the genomic integration spot. This linear DNA fragment was introduced in the cell by electroporation after one hour of inducing the λ-Red recombineering machinery with 15 mM arabinose. The cell suspension was then plated on LB plates containing the relevant antibiotics and a colony PCR was performed to confirm successful gene deletions. The antibiotics cassettes were removed by inducing the flippase with 50 µM L-rhamnose for at least four hours and removal was confirmed by a colony PCR. M2 was integrated into the genomic safe spot 9 (Bassalo et al., 2016) using a kanamycin marker and expressed under the control of a strong constitutive promoter (P_pgi-20_) and a medium strength RBS (RBS_C_). To create the formaldehyde sensor, the gene *rhmA* from *E. coli* which encodes for the HAL reactions was integrated into the genomic safe spot 2 (Bassalo et al., 2016) using a kanamycin marker under the control of P_pgi-20_ and RBS_C_. Strains used in this study are listed in **Table 1**.

**Table 1:** Strains used in this study.

| Strain name | Gene deletions | Gene insertion | Plasmid | Source | Used in figure |
| --- | --- | --- | --- | --- | --- |
| SerAux | $\Delta gcvTHP \Delta serA$ | none | none | This study | Suppl. Fig. 1 |
| SerAux | $\Delta gcvTHP \Delta serA$ | none | M1 | This study | Suppl. Fig. 1 |
| GFP sensor | $\Delta gcvTHP \Delta serA \Delta frmRAB$ | none | pTR47m4-GFP | This study | Fig. 3 |
| GFP sensor + M2 | $\Delta gcvTHP \Delta serA \Delta frmRAB$ | SS9:$P_{pgi-20}$:rbsC:ltaE | pTR47m4-GFP | This study | Fig. 3 |
| GFP sensor + M1 + M2 | $\Delta gcvTHP \Delta serA \Delta frmRAB$ | SS9:$P_{pgi-20}$:rbsC:ltaE | pTR47m4-GFP pM1 | This study | Fig. 3 |
| Formaldehyde sensor | $\Delta gcvTHP \Delta serA \Delta frmRAB \Delta asd$ | SS2:$P_{pgi-20}$:rbsC:rhmA | none | This study | Fig. 4 |
| Formaldehyde sensor + M1 | $\Delta gcvTHP \Delta serA \Delta frmRAB \Delta asd$ | SS2:$P_{pgi-20}$:rbsC:rhmA | pM1 | This study | Fig. 4 |
| Formaldehyde sensor + M1 + M2 | $\Delta gcvTHP \Delta serA \Delta frmRAB \Delta asd$ | SS2:$P_{pgi-20}$:rbsC:rhmA, SS9:$P_{pgi-20}$:rbsC:ltaE | pM1 | This study | Fig. 4, Fig. 5 |

### Synthetic-Operon construction

All genes were codon optimized for *E. coli* K-12, after which an 6xHis-tag was added to the N-terminal side. Subsequently, the genes were inserted into a cloning vector, which attaches RBS_C_ upstream of the gene (Zelcbuch et al., 2013). These cloning vectors were used to assemble synthetic operons via restriction and ligation using the protocol as described in (Wenk et al., 2018). The assembled operon was then cloned into an expression vector pZ (p15A origin of replication, streptomycin marker) under the control of P_pgi-20_ (Wenk et al., 2018). pTR47m4-GFP (Woolston et al., 2018) was used to construct the GFP sensor. Plasmids used in this study are listed in **Table 2**.

**Table 2:** Plasmids used in this study.

| Plasmid name | Genes | Source | GeneBank ID |
| --- | --- | --- | --- |
| pTR47m4-GFP |  | (Woolston et al., 2018) |  |
| M1 | p15A-ori; Strep<sup>R</sup>, $P_{pgi-20}$: $RBS_C$: *ftfL-fch-mtdA-glyA* | This study | *ftfL* Q83WS0<br>*fchA* Q49135<br>*mtdA* P55818<br>*glyA* P0A825 |

### Growth experiments

Prior to all growth experiments, the strains were streaked out from the glycerol stock on LB plates (supplemented with the respective antibiotic and 0.25 mM DAP in case of the formaldehyde sensor) and grown overnight at 37 °C. From these plates, precultures in relaxing liquid M9 medium (as indicated in **Table 3**) were inoculated for overnight growth at 37 °C and 220 rpm. The cells were harvested (8000 rpm, 3 min), washed three times with M9 minimal medium, and diluted to a final OD_600_ of 0.01 in the respective medium. In a Nunc 96-well microplates (Thermo Fisher Scientific), 150 µL of the culture was added to each well and the wells were covered with 50 µL of mineral oil (Sigma-Aldrich) to prevent evaporation. Growth experiments were performed in the BioTek Epoch2 plate reader (BioTek Instrument, USA), which measured the optical density after each kinetic cycle. A cycle consists of 12 shaking steps in which 60 seconds linear and 60 seconds orbital shaking were alternated (1 mm amplitude). The growth experiments that required fluorescence measurement were performed in a Tecan Infinite 200 Pro plate reader (Tecan, Switzerland). The OD_600_ was corrected to present the OD_600_cuvette_ by dividing the OD_600_plate_ by the correction factor 0.23. The green fluorescence was normalized by the OD_600_plate_. The mean of the technical duplicates is shown in the growth curves.

**Table 3:**
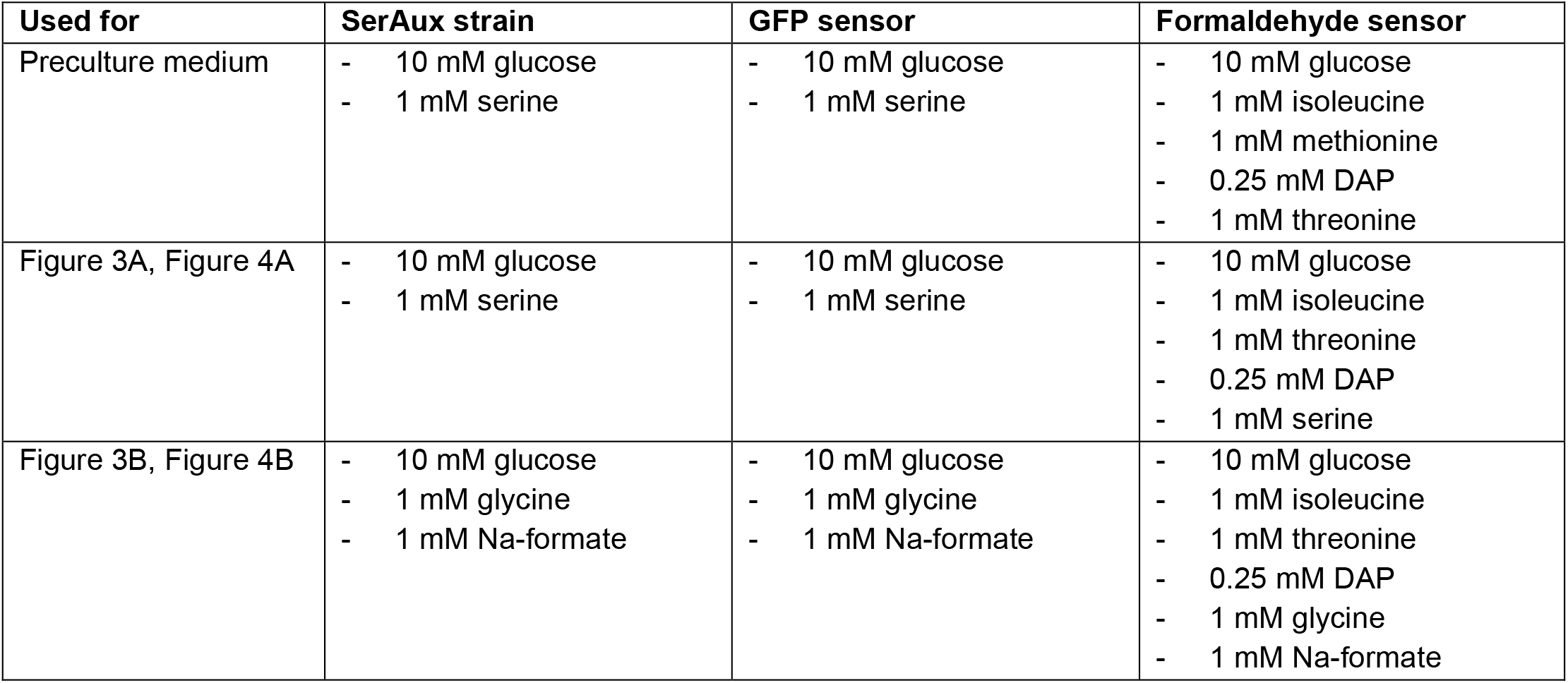
Composition of media for growth experiments.

### ^13^C labelling of proteinogenic amino acids

For stationary isotope tracing of proteinogenic amino acids, cells were cultured in 4 ml of M9 medium with all required carbon sources and supplements (10 mM glucose, 1 mM isoleucine, 1 mM threonine, 0.25 mM DAP, 1 mM glycine) as well as either labeled or unlabeled 1 mM Na-formate. After reaching the stationary phase, the cells were collected by centrifugation for 3 min at 8000 rpm. Biomass was hydrolyzed by incubation with 1 ml of 6N hydrochloric acid for a duration of 24 h at 95 °C. Samples were dried via heating at 95 °C and re-dissolved in 1 ml of ddH_2_O. Hydrolyzed amino acids were separated using ultra-performance liquid chromatography (Acquity, Waters) using a C18-reversed-phase column (Waters) as previously described in (Giavalisco et al., 2011). Mass spectra were acquired using an Exactive mass spectrometer (Thermo Fisher).

Data analysis was performed using Xcalibur (Thermo Fisher). Before analysis, amino acid standards (Sigma-Aldrich) were analyzed under the same conditions to determine typical retention times.

## Results

### The metabolic architecture of the Serine Shunt

We designed the Serine Shunt based on the formate assimilation reactions of the Serine Cycle reactions and the serine aldolase reaction of the Homoserine Cycle (Anthony, 2011; He et al., 2020). To facilitate the testing of the shunt, we divided it into two metabolic modules (**Figure 2**). Module 1 (M1) converts formate and glycine to serine and module 2 (M2) cleaves serine to formaldehyde and glycine. In M1, formate undergoes a series of enzymatic transformations catalysed by *Methylorubrum extorquens’* formate THF ligase (*Me*FtfL), methenyl-THF cyclohydrolase (*Me*Fch) and methylene-THF dehydrogenase (*Me*MtdA), consuming ATP and NADPH. The resulting methylene-THF then combines with glycine via *E. coli*’s serine hydroxymethyltransferase (*Ec*GlyA) to form serine. In M2, serine is cleaved into formaldehyde and glycine via the serine aldolase reaction catalysed by the threonine aldolase from *E. coli* (*Ec*LtaE). Notably, glycine acts as catalysing agent between both modules without being consumed by the pathway itself.

**Figure 2:**
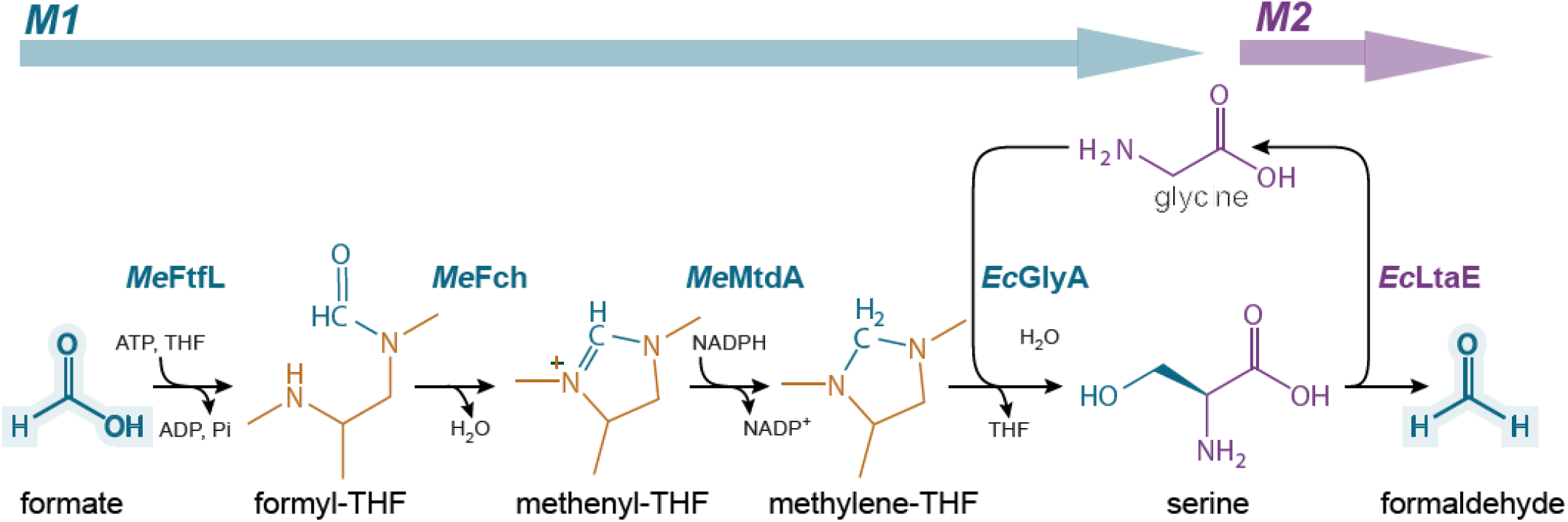
Molecular architecture of the Serine Shunt. The Serine Shunt can be divided into two modules: M1 (blue arrow + enzyme names) and M2 (purple arrow and enzyme names). In M1, Formate (blue) is attached to tetrahydrofolate (THF) (orange) forming a C1-THF intermediate, which is reduced to methylene-THF. The reduced C1 is subsequently transferred from THF onto glycine (purple), resulting in serine. In M2, serine gets cleaved, releasing the C1 as formaldehyde and recovering glycine. Glycine serves as catayst between M1 and M2. THF tetrahydrofolate, *Me*FtfL formate THF ligase, *Me*Fch methenyl-THF cyclohydrolase, *Me*MtdA methylene-THF dehydrogenase, *Ec*GlyA serine hydroxymethyltransferase, *Ec*LtaE threonine aldolase.

### Testing of pathway modules confirms *in vivo* activity of the Serine Shunt

While both modules of the Serine Shunt have been used in other metabolic context (He et al., 2020; Kim et al., 2020; Wenk et al., 2022), we assessed their individual activity first, before combining them into the complete shunt. To test M1, we introduced a plasmid expressing the module into a serine auxotrophic *E. coli* strain deleted in the phosphoglycerate dehydrogenase (Δ*serA*) and the glycine cleavage system (Δ*gcvTHP*) (SerAux strain) (**Supplementary Figure 1A**). While Δ*serA* creates a serine auxotrophy, which allows to select for formate conversion into serine, Δ*gcvTHP* prevents methylene-THF formation from glycine, ensuring dependency on formate assimilation. This strain was only capable of growing on glucose and glycine when we expressed M1 and supplemented formate (**Supplementary Figure 1B and 1C**).

Subsequently, we assessed whether *Ec*LtaE can produce formaldehyde from serine *in vivo* and thus, can be harnessed as M2 of the Serine Shunt. To this end, we deleted the formaldehyde detoxification system (Δ*frmRAB*) in the SerAux strain to prevent formaldehyde oxidation to formate. Subsequently, we introduced a formaldehyde-sensing plasmid (developed by Woolston al. (Woolston et al., 2018)) that allows to detect intracellular formaldehyde, by generating a GFP signal upon formaldehyde detection and referred to the resulting strain as GFP sensor (**Figure 3A**). To test M2 activity in this strain, we cultivated it on glucose and serine (**Figure 3B**). Only when *Ec*LtaE was overexpressed a GFP signal could be detected, confirming the enzyme’s ability to catalyse the conversion of serine into formaldehyde and its suitability as M2 of the Serine Shunt.

**Figure 3:**
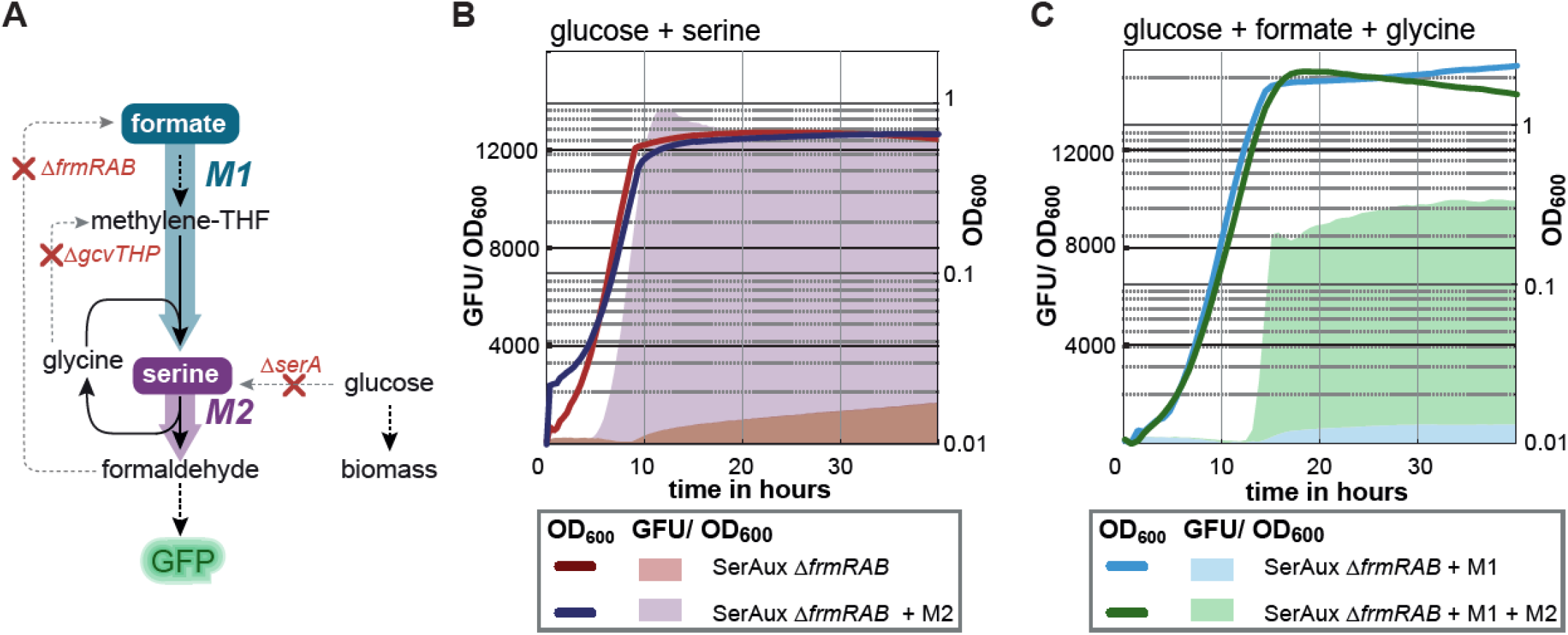
Testing of Serine Shunt modules in the GFP sensor. (A) Metabolic scheme of the GFP sensor. Gene deletions are indicated as red crosses. Multi-enzymatic reactions are shown as dotted arrows. As before, M1 is shown in blue colour and M2 in purple. (B) M2 activity was tested in the GFP sensor, cultivated on glucose as biomass precursor and serine as formaldehyde precursor. The green fluorescence normalized to the OD_600_ (GFU/ OD_600_) is shown on the left y-axis (shadow area) and the OD_600_ (curve) on the right one. Despite similar growth phenotypes between the GFP sensor with M2 overexpression (purple curve) and without (orange curve), only the strain overexpressing M2 produced a fluorescent signal (purple shadow). (C) The complete Serine Shunt was tested in the GFP sensor was cultivated on glucose as biomass source, glycine as catalysing reagent and formate as source of formaldehyde. Through M1 expression, formate could be converted to serine, releasing the strains’ auxotrophy (blue curve). Only upon expression of both modules green fluorescence could be measured, indicating the activity of the complete Serine Shunt. *frmRAB*, formaldehyde detoxification system, *gcvTHP* glycine cleavage system, *serA* serine hydroxymethyltransferase.

Having confirmed the individual activities of M1 and M2, we proceeded to evaluate their combined activity. To this end, we expressed both modules in the GFP sensor. We cultivated the resulting strain on glucose, glycine and formate and measured the GFP signal over time. While the strain was able to grow when M1 was expressed alone (release of the serine auxotrophy), only when both modules were expressed together a GFP signal was detected (**Figure 3C**). These results demonstrated for the first time the activity of the complete Serine Shunt.

### The Serine Shunt carries sufficient flux to support cell growth

While the activity of the Serine Shunt could be demonstrated using the GFP sensor, it’s essential to note that this sensor does not use formaldehyde as a source of biomass. To test if the Serine shunt can carry sufficient flux to support cell growth, we engineered the SerAux Δ*frmRAB* strain into a formaldehyde sensor which depends on formaldehyde assimilation for cellular growth. Leveraging the HOB sensor design from a previous studies (Schann et al., 2023), we deleted the aspartate-semialdehyde dehydrogenase (Δ*asd*) and overexpressed the HOB aldolase (HAL) in the SerAux Δ*frmRAB* strain. Δ*asd* rendered the strain auxotrophic to various essential amino acids, including methionine, while HAL enabled the assimilation of formaldehyde and pyruvate into the methionine precursor 4-hydroxy-2-oxobutanoate (HOB), thus releasing the strain’s auxotrophy in the presence of formaldehyde.

We cultivated the formaldehyde sensor, overexpressing M2, on glucose and serine to test whether serine cleavage by M2 could release the methionine auxotrophy. Only when M2 was overexpressed the strain’s growth could be rescued, thus, indicating that M2 can produce sufficient formaldehyde for biomass formation (**Figure 4B**). To test the complete Serine Shunt in a growth dependent context, we expressed both M1 and M2 in the formaldehyde sensor. The resulting strain was tested in a medium supplemented with glucose glycine and formate, requiring formate reduction to formaldehyde to release the methionine auxotrophy of the strain. As shown in **figure 4C** only when both modules were expressed, the strains growth was restored, indicating that the Serine Shunt is active *in vivo* and can convert formate into formaldehyde in an amount sufficient for biomass formation.

**Figure 4:**
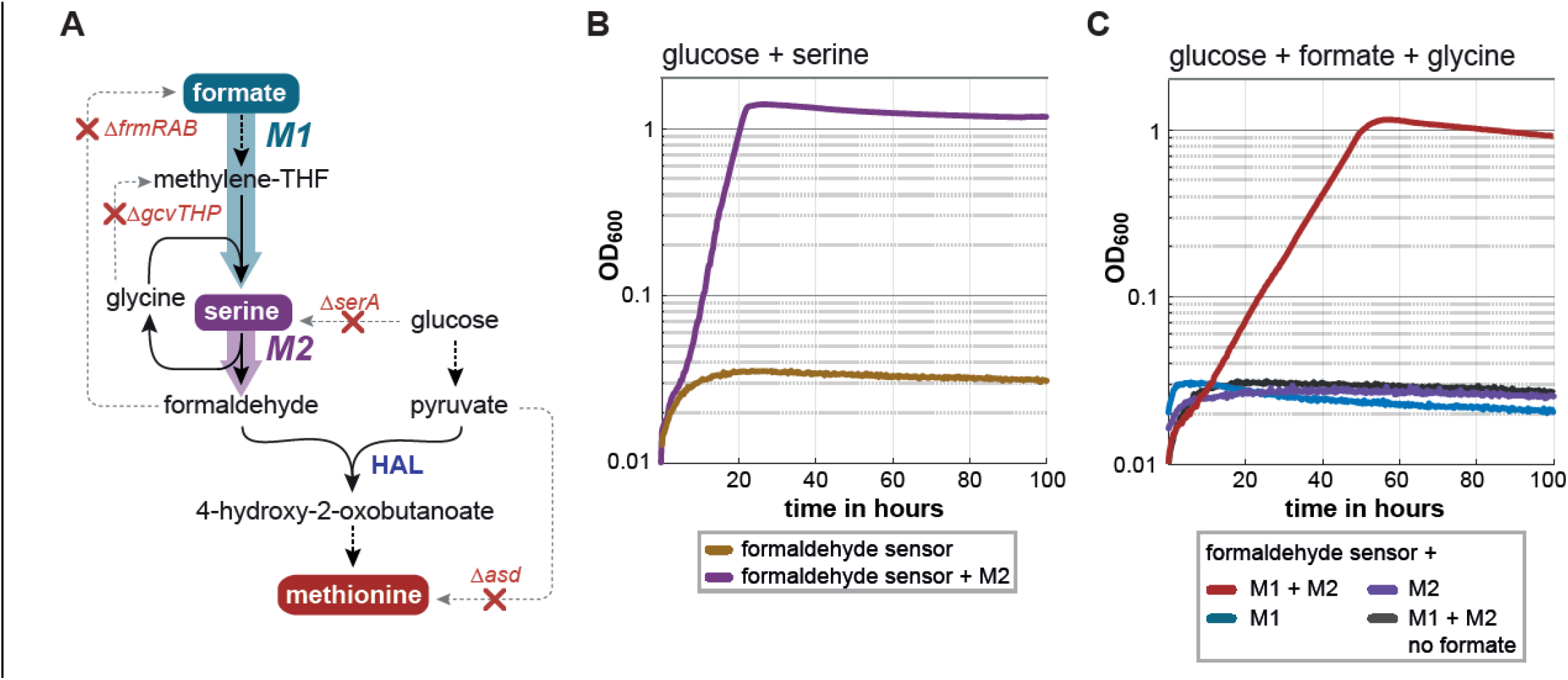
The Serine Shunt supports formate dependent growth. (A) The metabolic scheme of the formaldehyde sensor which was used to test the complete Serine Shunt. The methionine auxotrophy is shown as red circle. Gene deletions are visualized with a red cross. Multi-enzymatic reactions are indicated by dotted arrows. M1 is visualized in blue colour, whereas M2 is shown in purple. (B) The formaldehyde sensor was used to test M2 activity. M2 overexpression (purple curve) is sufficient to enable the strains growth on glucose (pyruvate precursor) and serine (formaldehyde precursor). (C) Combining both modules in the formaldehyde sensor (red curve) enables growth on glucose (pyruvate precursor), formate (formaldehyde precursor) and glycine (catalysing agent). *asd* aspartate-semialdehyde dehydrogenase, *frmRAB*, formaldehyde detoxification system, *gcvTHP* glycine cleavage system, *serA* serine hydroxymethyltransferase, HAL HOB aldolase.

### Carbon tracing confirms formate conversion to formaldehyde via the Serine Shunt

To confirm that formaldehyde is produced from formate via the Serine Shunt, we used ^13^C-labellling to track the metabolic incorporation of the formate-derived carbon in the formaldehyde sensor. In theory, the carbon from formate is incorporated into serine via M1 and cleaved into formaldehyde by M2. Thereby, glycine serves only as catalyser between the modules and doesn’t mix with the formate-derived carbon. Following this, serine should be labelled once, while glycine remains unlabelled. Formaldehyde derived from serine cleavage condenses with pyruvate to HOB, which gets converted into homocysteine. Homocysteine condenses with methylene-THF coming from formate, forming methionine. As methylene-THF is derived directly from formate, methionine should be labelled twice **(Figure 5A)**.

We cultivated the formaldehyde sensor expressing both modules on glucose, glycine and either non-labelled formate or ^13^C-labelled formate **(Figure 5B)**. Then, we extracted the proteinogenic amino acids and analysed their labelling pattern via liquid chromatography mass spectrometry (LC-MS). When ^13^C formate was supplied in the medium, all amino acids were labelled as expected. Serine was labelled once, confirming M1 activity. Methionine showed two labelled carbons, confirming the activity of the complete Serine Shunt. Thus, the ^13^C-labelling results unequivocally confirm the *in vivo* activity of the Serine Shunt.

**Figure 5:**
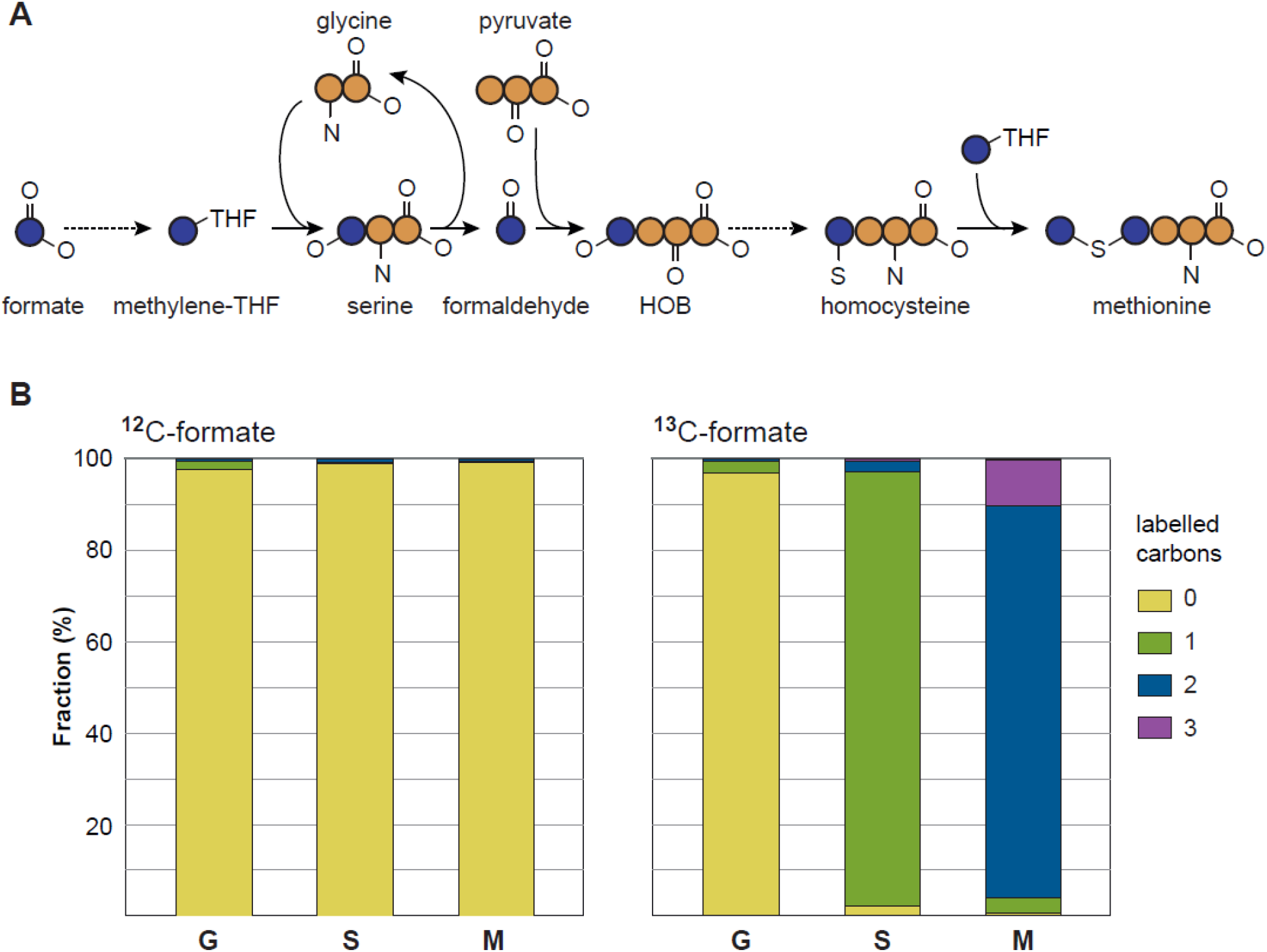
13C labelling confirms *in vivo* activity of the Serine Shunt. (A) Expected labeling of proteinogenic amino acids upon cultivation on ^13^C-formate. The carbon derived from formate is colored blue, whereas carbons from all other sources are orange. Multi-enzymatic reactions are indicated by dotted arrows. (B) The strain fed with ^12^C-formate shows no labelling in glycine (G), serine (S) and methionine (M). (A) When ^13^C-formate was used as formaldehyde precursor, serine is labelled once (green) and methionine is labelled twice (blue). HOB 4-hydroxy-2-oxobutanoate.

## Discussion

The quest for a sustainable, CO_2_-based bioeconomy has resulted in massive research efforts in recent years (Bachleitner et al., 2023). While CO_2_ and other C1 gases like carbon monoxide and methane could be envisioned as substrates for bacterial cultivation, liquid feedstocks like methanol and formate present multiple advantages ranging from their relative ease of handling to better mass transfer rates (Cotton et al., 2020). Of the liquid substrates, formate can be produced with high efficiencies in a one-step reaction from CO_2_. Thus, formate is considered a prime candidate feedstock for a CO_2_-based bioeconomy (Claassens et al., 2019). Until now, a formate bioeconomy has been postulated and several microbes have been engineered to use formate as a feedstock (Bang et al., 2020; Bruinsma et al., 2023; Bysani et al., 2024; Claassens et al., 2020; Kim et al., 2020; Turlin et al., 2022; Wenk et al., 2022).

An undesired fact that limits the potential of a formate bioeconomy is the limited number of formate fixing enzymes and the associated small number of metabolic pathways that can be designed from them (Bar-Even, 2016). A “metabolic bridge” or “metabolic bypass” (Orsi et al., 2022) that connects formate to formaldehyde would vastly increase the potential of a formate bioeconomy as it would allow to convert methylotrophs into formatotrophs. As enzymatic formate reduction to formaldehyde does not seem to exist in nature, new-to-nature metabolic pathways for this conversion need to be established. This could be achieved by novel enzyme chemistries as suggested for the formyl-CoA and the formyl phosphate route as detailed in (Erb et al., 2017) or by metabolic rewiring as proposed for the Serine Shunt. The latter has the advantage of using established enzyme activities that do not require substantial enzyme engineering. In this work, we present the Serine Shunt as a promising alternative to the previously proposed formate reduction routes. The Serine Shunt uses the formate assimilation module of the natural Serine Cycle to convert formate and glycine into serine and the formaldehyde assimilation module of the Homoserine Cycle alas in reverse direction for cleaving serine to formaldehyde and glycine. In the Serine Shunt, glycine can be considered as a “catalyst” as it is essential to catalyze the reactions but is not consumed by the pathway. While longer in terms of enzymatic reactions, the Serine Shunt resembles previously proposed formate reduction routes in its structure and consumes the same amount of ATP and NADPH as the most energy efficient formyl phosphate route (Bar-Even, 2016; Nattermann et al., 2023). Furthermore, it only uses native enzymes allowing immediate implementation without enzyme engineering.

By dividing the pathway into two metabolic modules and coupling module activity to cell growth (growth-coupled selection), we showed that the individual modules are sufficiently active to support cell growth. Our results demonstrated for the first time that the serine aldolase used for formaldehyde assimilation in the synthetic Homoserine cycle (He et al., 2020) can efficiently cleave serine to glycine and formaldehyde *in vivo*. Next, we combined both modules into the complete Serine Shunt and achieved sufficient flux to generate essential cellular building blocks from formate. These results demonstrate that the Serine Shunt is a promising route to establish cell growth via formate reduction.

While in this work, only parts of cellular biomass were derived from formate, in future studies, the Serine Shunt could be tested in methylotrophs that grow via formaldehyde assimilation pathways like the RuMP cycle. Here, the Serine Shunt could serve as metabolic route that generates all biomass from formate paving the way to a sustainable, CO_2_-based formate bioeconomy.

## Acknowledgements

This work was supported by the German Ministry of Education and Research Grant 031B0850B (MetAFor) and the Max Planck Society. The authors are grateful to Arren Bar-Even who inspired the project and was a brilliant mentor to all of us. The authors thank Saleh Alseekh for performing the HPLC-MS measurements and Enrico Orsi for critical reading of the manuscript and helpful suggestions.

## Author contributions

S.W. conceptualized the project and supervised the research.

S.W. and K.S. designed the experiments

K.S. conducted the experiments.

S.W. and K.S. analyzed the data and conceived the manuscript.

## Data availability

Additional information on the experimental setup as well as detailed results are available from the corresponding author upon request. Any strains and plasmids generated during this study are available upon completing a Materials Transfer Agreement.

