## Supplementary Figure 1 for "The Serine Shunt enables formate conversion to formaldehyde *in vivo*"

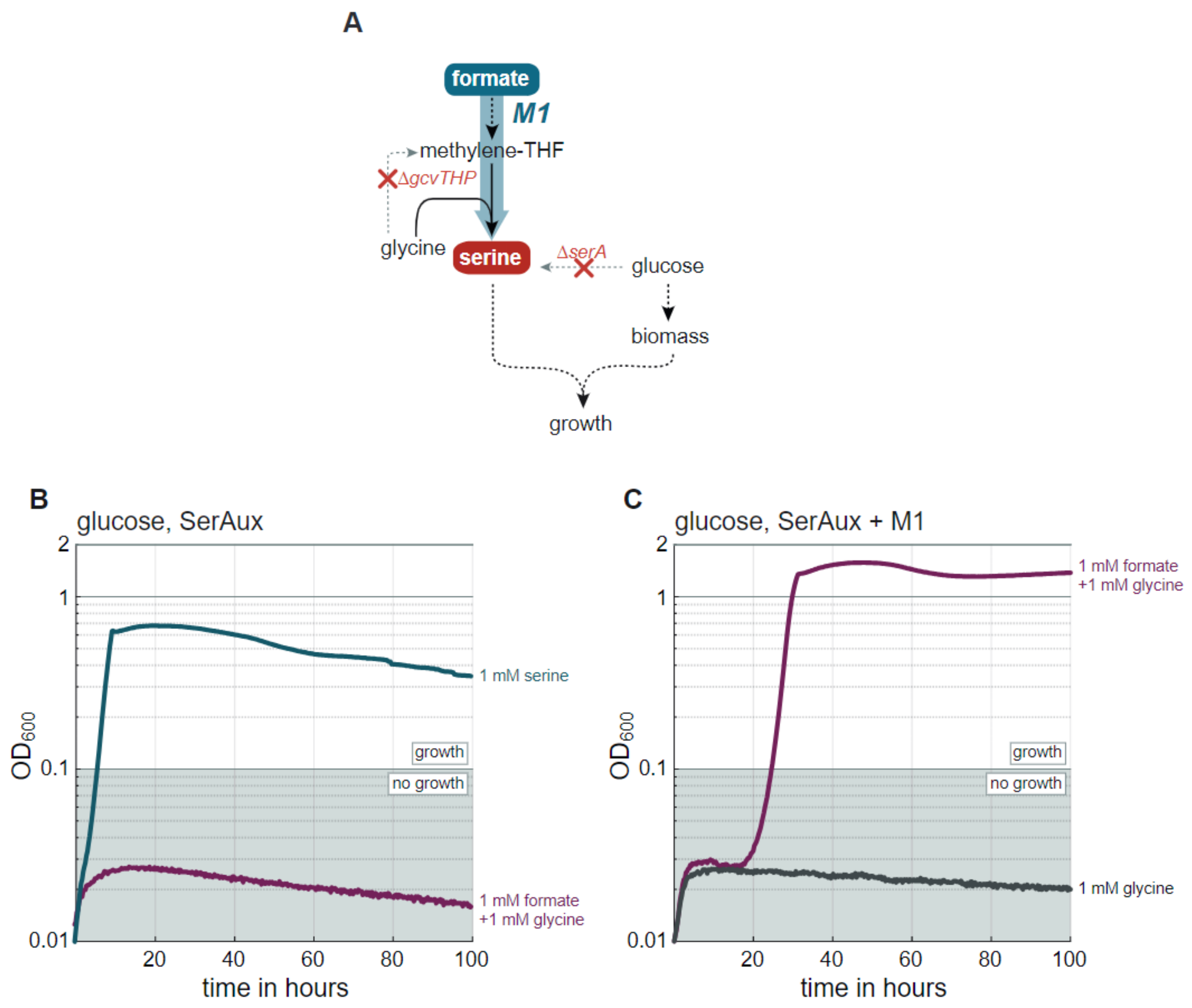


**Supplementary Figure 1: Testing M1 activity in SerAux strain**

(A) Metabolic scheme of the SerAux sensor. Gene deletions are indicated as red crosses. Multi-enzymatic reactions are shown as dotted arrows. M1 is shown in blue colour. (B) The SerAux strain is only capable to grow on glucose when serine is supplemented additionally (blue curve). (C) M1 activity was tested in the SerAux sensor, cultivated on glucose as biomass precursor, formate as formaldehyde precursor and glycine as catalysing agent. and serine as formaldehyde precursor. Through M1 expression, formate could be converted to serine, releasing the strains' auxotrophy (purple curve). *gcvTHP* glycine cleavage system, *serA* serine hydroxymethyltransferase.
